# Viscoelastic recovery time of chondrocytes from monolayer and alginate cultures

**DOI:** 10.64898/2026.02.16.706204

**Authors:** Michael Neubauer, Priyanka Brahmachary, Ronald K. June, Stephan Warnat

**Author notes:** Corresponding Author: Stephan Warnat.

## Abstract

This study uses a previously reported 3D-printed variable-height flow cell to investigate the viscoelastic properties of chondrocytes from 2D monolayer and 3D alginate cultures. It was hypothesized that chondrocytes could be distinguished by phenotype associated with their culture environment using viscoelastic recovery time, owing to variation in the pericellular matrix (PCM) produced by chondrocytes from different culture methods. The PCM surrounding the chondrocytes was imaged with confocal microscopy during applied deformation and subsequent recovery. The projected cell area was fitted with a Burgers mechanical model to extract the viscoelastic recovery time. No difference between bovine and primary OA cells from monolayer cultures was observed. However, a statistically significant difference in recovery time was observed between cells from monolayer and alginate cultures in both the bovine (31 s vs. 13 s) and primary OA (34 s vs. 13 s) groups. This work shows that viscoelastic recovery time is influenced by the culture method used for chondrocytes and further demonstrates the role of the PCM as a mechanical protector of chondrocytes.

## Introduction

Osteoarthritis (OA) is the most common degenerative joint disease, impacting more than 500 million people worldwide. Of these cases, over 300 million involve the knee joint, a particularly vulnerable site due to its high load-bearing function [1]. OA leads to the breakdown of the articular cartilage that lines bone surfaces within joints, which results in pain, stiffness, and reduced mobility over time [2]. Articular cartilage primarily comprises an extracellular matrix (ECM), a supportive and flexible network that provides structural integrity and shock absorption. Chondrocytes, the only resident cells within the cartilage, are crucial for cartilage health, functioning, and sustaining this matrix. [3]. Surrounding each chondrocyte is a thin, 2–4 µm layer of pericellular matrix (PCM), forming what is known as a chondron. The PCM acts as a buffer between chondrocytes and the ECM, facilitating essential cell functions, including mechanotransduction, in which cells convert mechanical stimuli into biochemical signals [4-6]. The PCM plays a vital role in protecting chondrocytes from mechanical stress. Through mechanotransduction, chondrocytes can sense and respond to mechanical forces by adjusting their gene expression and metabolic activity, which is critical for maintaining cartilage homeostasis and potentially slowing OA progression [7].

Chondrocytes cultured in a monolayer environment often undergo dedifferentiation, shifting from their native spherical shape *in vivo* to a flattened, elongated 2D morphology [8]. Additionally, monolayer cultures do not produce chondrocytes with well-defined PCM layers. Evidence suggests that the PCM is crucial for maintaining chondrocyte identity, as studies have shown that isolated chondrons (chondrocytes with intact PCM) generate a more cartilage-like ECM compared to monolayer-cultured chondrocytes, which produce a less authentic cartilage matrix. [9, 10]. To promote a more *in vivo*-like phenotype, chondrocytes are often cultured within millimeter-scale alginate beads or hydrogels [11-13]. This 3D environment promotes the cells’ retention of their rounded morphology, closely resembling the morphology of *in vivo* chondrocytes. However, chondrocytes grown in alginate beads can either lack an endogenous PCM or exhibit a reduced PCM stiffness [14-18]. Ideally, chondrocytes or chondrons used in experimentation should mimic the *in vivo* phenotype and possess biophysical properties similar to those of native cells, so that results can be directly applied to cells in their natural state and environment. More evaluation of culture methods is required.

The PCM directly transfers the load from the ECM to the chondrocyte [19-21]. Chondrocytes have been suspended in high-stiffness agarose gel to study *in vitro* cellular response to mechanical loading [22-25]. Viscoelastic mechanical properties of chondrocytes have been previously measured with micropipette aspiration (MPA) [26-32] and atomic force microscopy (AFM) [33-36], but typically these cells have been cultured in 2D monolayers. The PCM of chondrons was measured with AFM [13, 37] and MPA [38], and the elastic modulus of the PCM was found to be up to 10 times higher than that of the chondrocyte [5, 39, 40]. The effect of OA on the chondrocyte PCM remains controversial, as studies using MPA have reported increased chondrocyte stiffness with the progression of OA [26-28]. In contrast, studies with AFM [35, 41] and confined compression [42] showed the opposite. A more thorough explanation of the chondrocyte isolation and culture in these studies could highlight the discrepancy in the results.

This work used a previously reported 3D-printed variable-height microfluidic device for viscoelastic characterization of bovine and primary OA chondrocyte cells from 2D monolayer and 3D alginate cultures. Bovine chondrocytes were used as a healthy control for primary OA cells, and a standard monolayer culture was a control for a recently reported alginate culture method [43]. Both monolayer and alginate cultures include sodium L-ascorbate, which promotes increased collagen VI which is a major component of the PCM. The alginate culture was previously shown to produce a more robust PCM shell than cells cultured in monolayers. It was hypothesized that the viscoelastic properties of cells from monolayer and alginate cultures would be distinct, given the increased stiffness of PCM relative to chondrocytes. Cells of each type, from each culture method, were loaded into the device and subjected to mechanical compression and relaxation while imaged using confocal laser scanning microscopy (CLSM). The projected area of the cells was fit to a Burgers four-element mechanical model, and the viscoelastic recovery time was compared between the groups of N=3 donors each. Quantifying viscoelastic recovery time offers an additional perspective on how culture‐dependent differences in the PCM may influence whole‐chondron mechanical behavior, complementing prior single‐cell and PCM‐focused mechanical studies.

## Methods

### Flow cell and CLSM imaging

Previous work [44] describes the fabrication of the variable-height 3D-printed device used in this study. Briefly, a channel structure was 3D-printed directly on a No. 1.5 glass coverslip treated with bind silane, as in [45]. A 3 mm x 4 mm piece of glass was placed in a cutout in the 3D-printed channel and sealed with silicone. Another 3D-printed structure, the control chamber, encloses the area above the glass piece, and a custom pressure controller controls the pressure inside. The height of the fluidic channel between the glass coverslip and the top glass piece is controlled by adjusting the pressure of the control chamber. For each experiment, cells are injected through tubing and flown into the variable height region, where they are compressed between the two pieces of glass and imaged with a confocal microscope (Stellaris DMi8, Leica, USA) using a 20x objective (HC PL APO 20x/0.75 CS2, dry objective, Leica, USA). The application and removal of cell compression by the movable glass piece occur quickly, less than 1 s, although the exact displacement rate was not measured. The compression time for each experiment was 30 s, and cell recovery was monitored for an additional 180 s.

### Chondrocyte Harvest and Culture

Four different groups of chondrocyte cells were used for experimentation: (i) bovine chondrocytes from monolayer cultures, (ii) bovine-alginate, (iii) primary OA-monolayer, and (iv) primary OA-alginate, as shown in Fig. 1. Bovine cells were used as a healthy control. Three donors (N=3) comprised each group. Human Primary Chondrocytes (HPC) were obtained from discarded arthroplasty tissue from Stage-IV osteoarthritis patients undergoing total joint replacement under IRB approval. Bovine chondrocytes were harvested from knee joints of 18-22-month-old cattle obtained from a local abattoir. Briefly, tissue was digested with Collagenase Type I (2 mg/mL) (Gibco, Waltham, MA, USA) for 14 h at 37ºC using a previously established protocol [46]. Isolated chondrocytes were cultured for 10 days in Dulbecco’s Modified Eagle’s medium (DMEM) (Gibco, Waltham, MA, USA) supplemented with Fetal Bovine Serum (FBS) (10% v/v) (Bio-Techne, Minneapolis, MN, USA), penicillin (10,000 I.U./mL), and streptomycin (10,000 µg/mL) (Sigma, St. Louis, MO, USA) (hereby referred to as Complete Media) in 5% CO_2_ at 37ºC and first passage population was used for experiments.

**Fig. 1.**
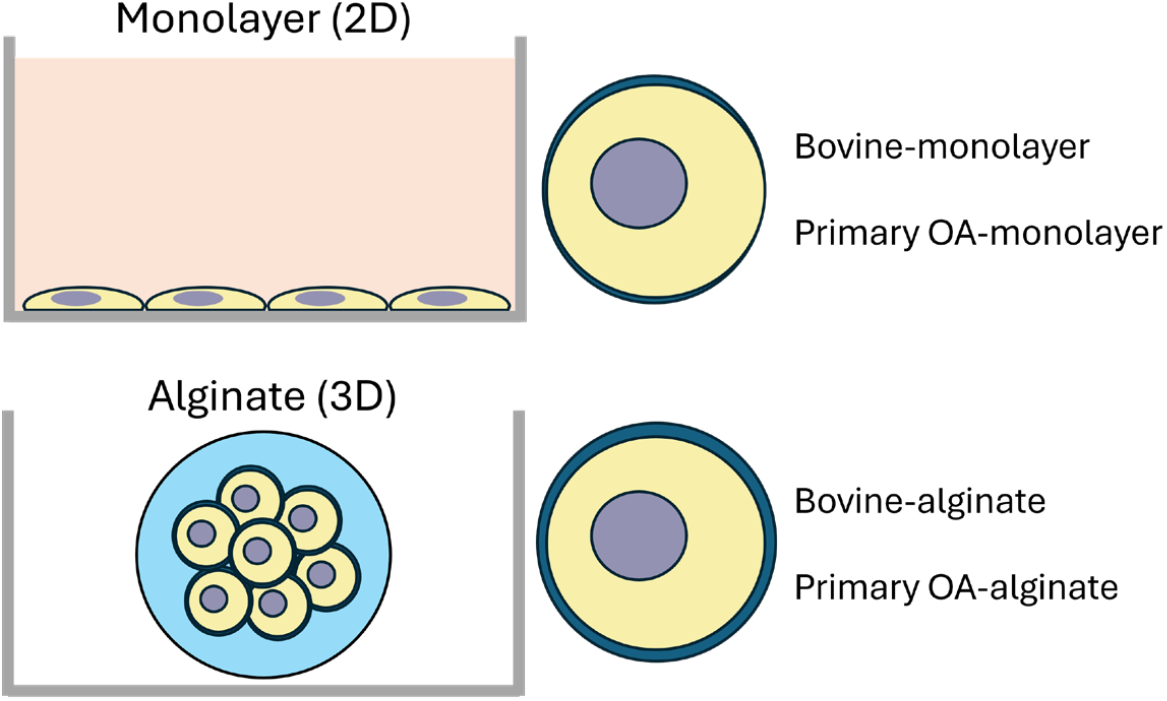
Monolayer vs alginate culture. Both cultures contain the addition of sodium L-ascorbate. Bovine and primary OA cells are produced from each culture methods, resulting in four different populations of chondrocytes for experimentation.

For monolayer studies, chondrocytes were cell cycle synchronized by 12 h starvation followed by culturing in Complete media and incubated in 5% CO_2_ at 37ºC for 7 days with 50 μg/mL sodium L-ascorbate (Sigma), and the media was changed every other day. For alginate encapsulation, cells that were cell-cycle synchronized were resuspended at a density of 4×10^6 cells/mL in sterile-filtered 1.2%, w/v, sodium alginate (Sigma). The cell suspension was then slowly dispensed through a 22-gauge needle in a dropwise fashion into a 102 mM CaCl_2_ solution. After near-instantaneous gelation, the alginate beads were allowed to crosslink for 15 min. The beads were then washed 4 times with 150 mM NaCl and once with Complete media before being cultured with complete media containing 50 μg/mL sodium L-ascorbate in 5% CO_2_ at 37ºC for 7 days, following which the alginate beads were dissolved with an EDTA-citrate buffer (150 mM NaCl in 55 mM sodium citrate with 50 mM EDTA, pH6.8). Released cells were then stained with Calcein AM and Wheat Germ Agglutinin.

### Calcein AM and Wheat Germ Agglutinin Staining

1×10^6 cells/ml monolayer and alginate-released cells that were previously cultured with 50 μg/mL sodium L-ascorbate for 7 days were resuspended in Phosphate Buffer Saline (PBS, 1X) and Calcein Blue AM Viability Dye (Invitrogen) was added at a final concentration of 5 µM. Cells were incubated in 5% CO_2_ at 37ºC in the dark for 30 min. Cells were washed twice with 1X PBS and incubated with 5 µM final concentration of Wheat Germ Agglutinin, Alexa Fluor 594 conjugate (Life Technologies Corporation) in 5% CO_2_ at 37ºC in the dark for 10 min. Cells were washed with 1X PBS and kept ready for visualization. The Wheat Germ Agglutinin was used to label the cell membrane and visualize the cells by confocal microscopy.

### Data analysis

Each Z-stack of images in an experiment’s time series is first converted into a single image using a max-intensity projection (MIP) in Imaris (Oxford Instruments, USA). The time series is imported into MATLAB (MathWorks, USA), and each frame is processed by a custom Sobel edge-detection method to extract the border of each cell in the field-of-view (FOV). The projected area of each cell is recorded and then converted to linear strain.

A four-element Burgers mechanical model fits the time-dependent linear strain, shown in Equation 1. The Burgers model consists of four elements: a spring in series with a parallel spring-dashpot, in series with another dashpot. The time-dependent strain, *ε*(*t*), is equal to the sum of the magnitudes of elastic, viscoelastic, and viscous strains. The viscoelastic portion of the recovery is a decaying exponential with the time constant, τ, which represents the viscoelastic recovery time. Previous work has demonstrated that the interior cell stain can be used to track cell recovery and find the viscoelastic recovery time [44], but in this study, the cell membrane stain is used to define the cells’ projected area.

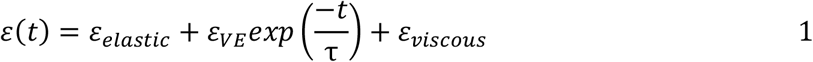

The projected area of each chondron, defined as the PCM boundary, is used to fit the recovery to the Burgers model, as illustrated in Fig. 2. The projected area is first converted to linear strain, but the general shape of the recovery is the same.

**Fig. 2.**
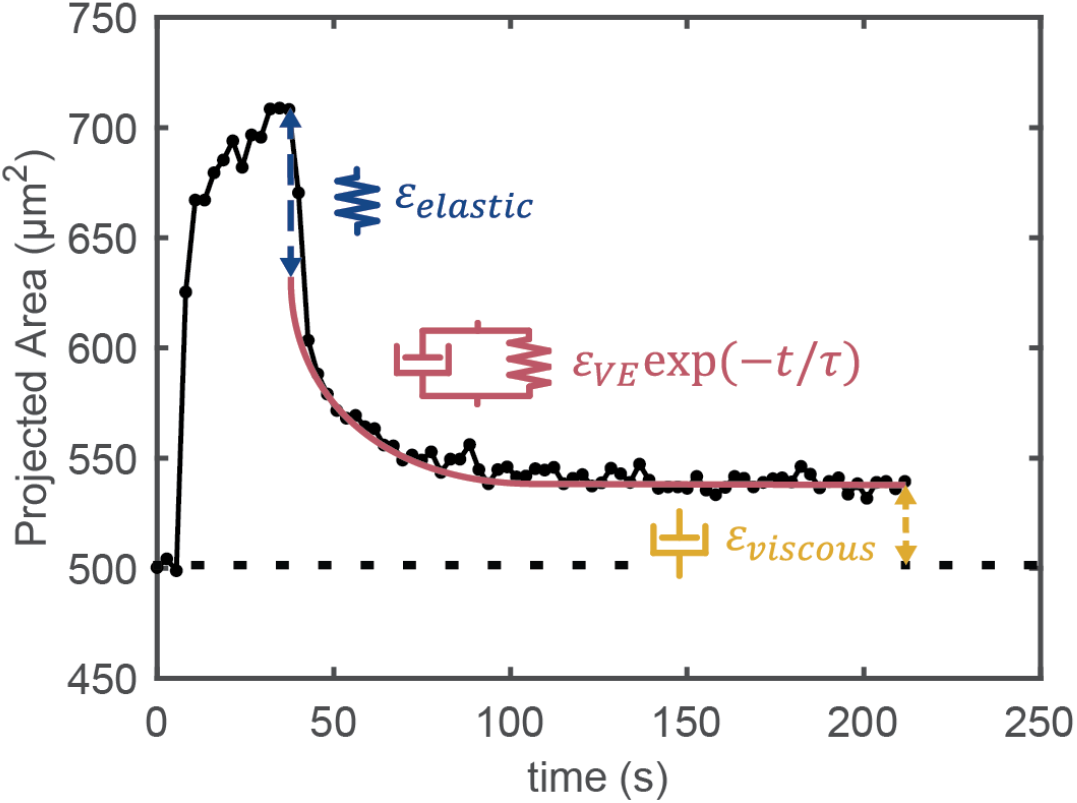
Representative data for the projected area of a chondron during recovery from an applied strain is fit to each mechanical element from a Burgers model. The chondrons have a nearly instantaneous elastic recovery, followed by an exponentially decaying viscoelastic recovery, characterized by the recovery time, τ. Any permanent deformation is accounted for by the dashpot element, representing the viscous strain.

## Results and discussion

### Cell Appearance

Bovine chondrocytes from monolayer and alginate cultures are shown in Fig. 3. The blue viability dye and red wheat germ agglutin (cell membrane) dye channels are overlaid in the first row and viewed separately in rows two and three, respectively. The two cells shown from each culture are representative of the cells analyzed in a recovery experiment. The PCM surrounding bovine chondrocytes is consistent between the monolayer and alginate cultures. However, there is some heterogeneity within cell populations, as shown in Fig. 4, which depicts a group of bovine monolayer cultures. Here, differences are apparent in chondrocyte size, shape, viability, and PCM thickness and uniformity. This image is representative of the heterogeneity observed in all cell populations in this work.

**Fig. 3.**
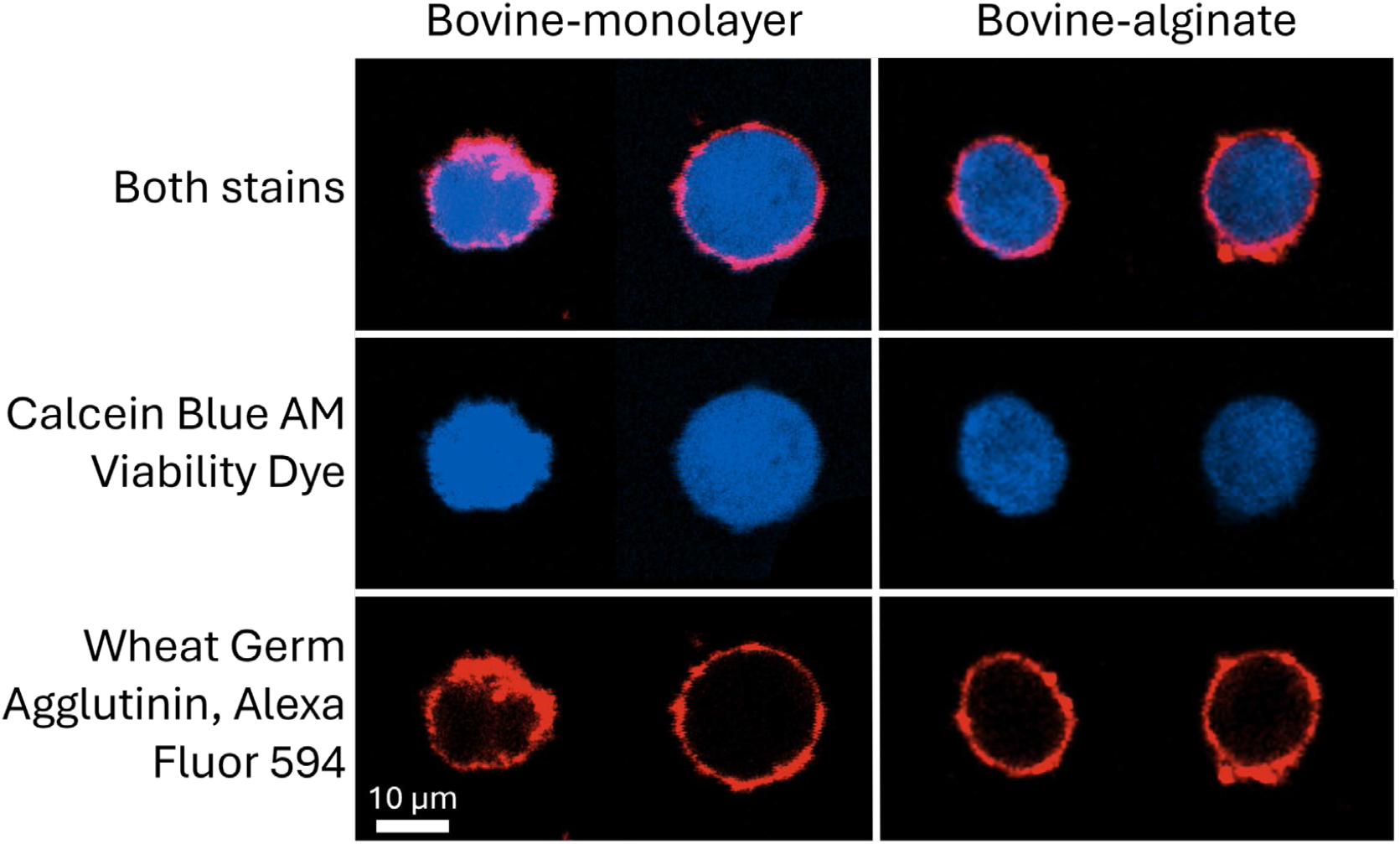
Confocal images (single slice from a Z-stack) of bovine chondrocytes from monolayer (left column) and alginate (right column) cultures prior to compression experiments. The scale bar in the bottom left corner is true for all images in this figure. The calcein blue AM viability dye stains the chondrocyte (center row) while the wheat germ agglutin stains the cell membrane of the chondrocyte (bottom row). The top row shows the two stains overlaid. The two cells from each culture are representative of the cells analyzed during a recovery experiment. In general, the PCM was visually similar between the monolayer and alginate cultured cells.

**Fig. 4.**
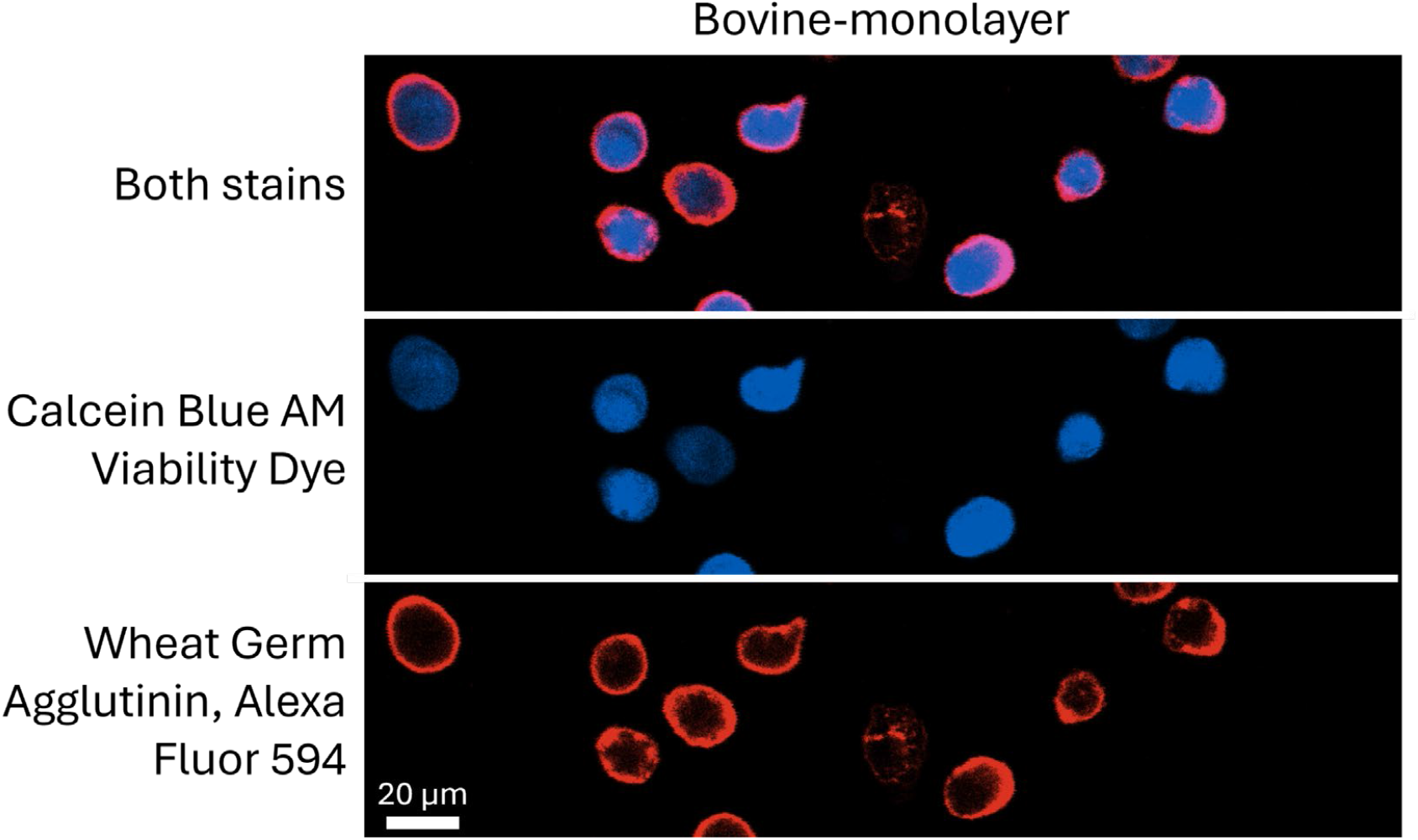
Larger field-of-view confocal images of bovine chondrocytes from a monolayer culture prior to compression experiments. The scale bar in the bottom left corner is true for all images in this figure. The calcein blue AM viability dye stains the chondrocyte (center row) while the wheat germ agglutin stains the cell membrane the chondrocyte (bottom row). The top row shows the two stains overlaid. This image is representative of the diversity of the chondron appearance. There are brighter and dimmer cells from the viability stain and the PCM thickness and uniformity varies from cell-to-cell. Note that this is a single Z-slice at one height and the cells shown are not all in the same plane. The apparent thickness of the PCM is influenced by the curvature of the cell within focus in the 0.68 µm thick Z-slice.

Similarly, representative primary OA chondrocytes and surrounding PCM from monolayer and alginate cultures are displayed in Fig. 5. Unlike bovine groups, the PCM thickness appears to be increased and more uniform in primary OA-alginate cultured chondrocytes compared to monolayer cultures.

**Fig. 5.**
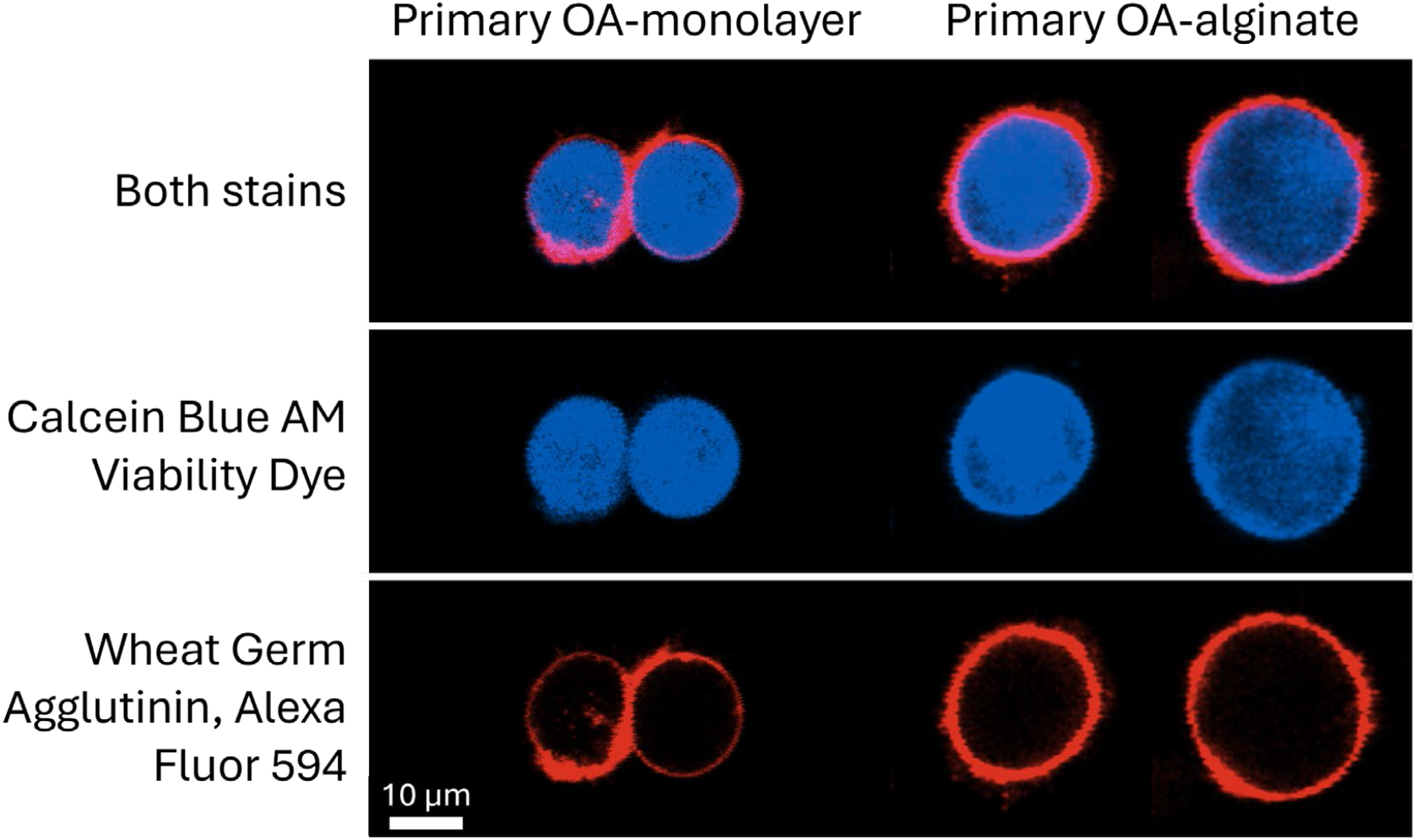
Confocal images (single slice from a Z-stack) of primary OA chondrocytes from monolayer (left column) and alginate (right column) cultures. The scale bar in the bottom left corner is true for all images in this figure. The calcein blue AM viability dye stains the chondrocyte (center row) while the wheat germ agglutin stains the cell membrane of the chondrocyte (bottom row). The top row shows the two stains overlaid. The two cells from each culture are representative of the cells analyzed during a recovery experiment. In general, the PCM produced from the alginate cultures appeared thicker and covered the chondrocytes more uniformly.

A thorough investigation of the PCM thickness and uniformity for all cell populations used in this work was not conducted. Still, it could be of interest to further characterize the culture methods. Previously, chondrocyte PCM thickness has been measured with optical microscopy [13, 47], confocal microscopy [48], and transmission electron microscopy [49]. It has also been reported that OA increased PCM thickness by up to 50% [27, 47]. The confocal microscope used in this work would be capable of measuring PCM thickness. However, the image acquisition mode and Z-slice thickness used in this work were selected for speed and are not ideal for this task.

### Viscoelastic Recovery Time

The permanent strain as a function of total applied strain for bovine and primary OA chondrons from monolayer cultures is shown in Fig. 6(a). In general, larger permanent strains result from higher applied strains. The height of the channel defines the strain of all the cells under the movable glass plate, so the initial diameter of the cells causes the variability of the applied strain, seen in Fig. 6(a-b). A previous study using the same device and general method found a similar result, but the data were from chondrocytes without a stained PCM [44]. Fig. 6(b) shows the viscoelastic recovery time of the same chondrons, and no dependence on total applied strain is observed.

**Fig. 6.**
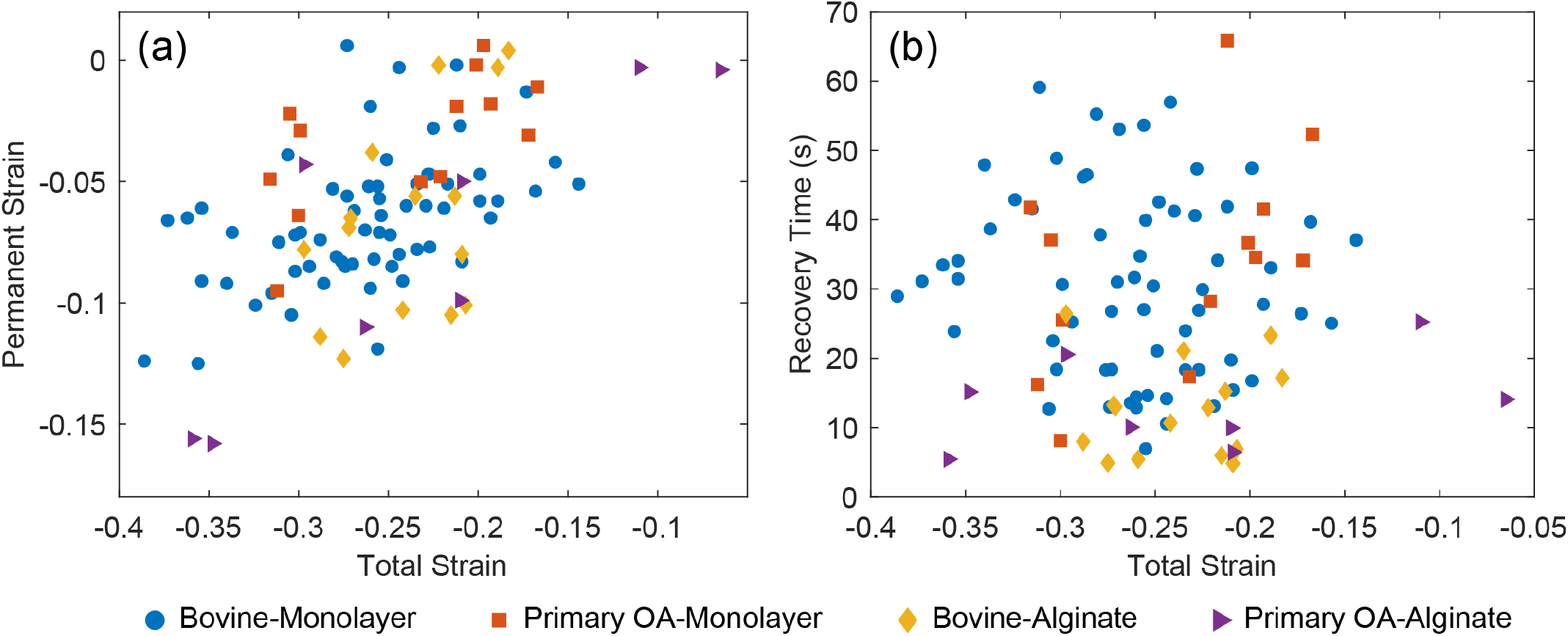
Permanent strain and recovery times for bovine and primary OA chondrons from monolayer and ate cultures. (a) Total strain vs permanent strain. There appears to be a rough correlation, the higher the ed strain, the greater the magnitude of the permanent strain. (b) the viscoelastic recovery time as aion of total applied strain. No relationship is apparent.

A comparison of the recovery times for each group of chondrons is shown in Fig. 7. The recovery time for bovine (mean=31 s, standard deviation (std)=13 s, n=64) and primary OA (mean=34 s, std= 15 s, n=13) chondrons from monolayer cultures showed no statistically significant difference. Similarly, the recovery time for bovine (mean=13 s, std = 7 s, n=15) and primary OA (mean=13 s, std = 7 s, n=8) chondrons was not deemed different. While no difference in recovery time was found between bovine and primary OA chondrons from the same culture method, chondrons compared between culture methods always resulted in a statistically significant difference, regardless of bovine or primary OA chondrons. Bovine chondrons showed a decreased recovery time from alginate cultures compared to monolayer cultures. The same was true for Primary OA chondrons. Welch’s unequal‐variance two‐sample t‐test (two‐tailed) to compare independent groups.

**Fig. 7.**
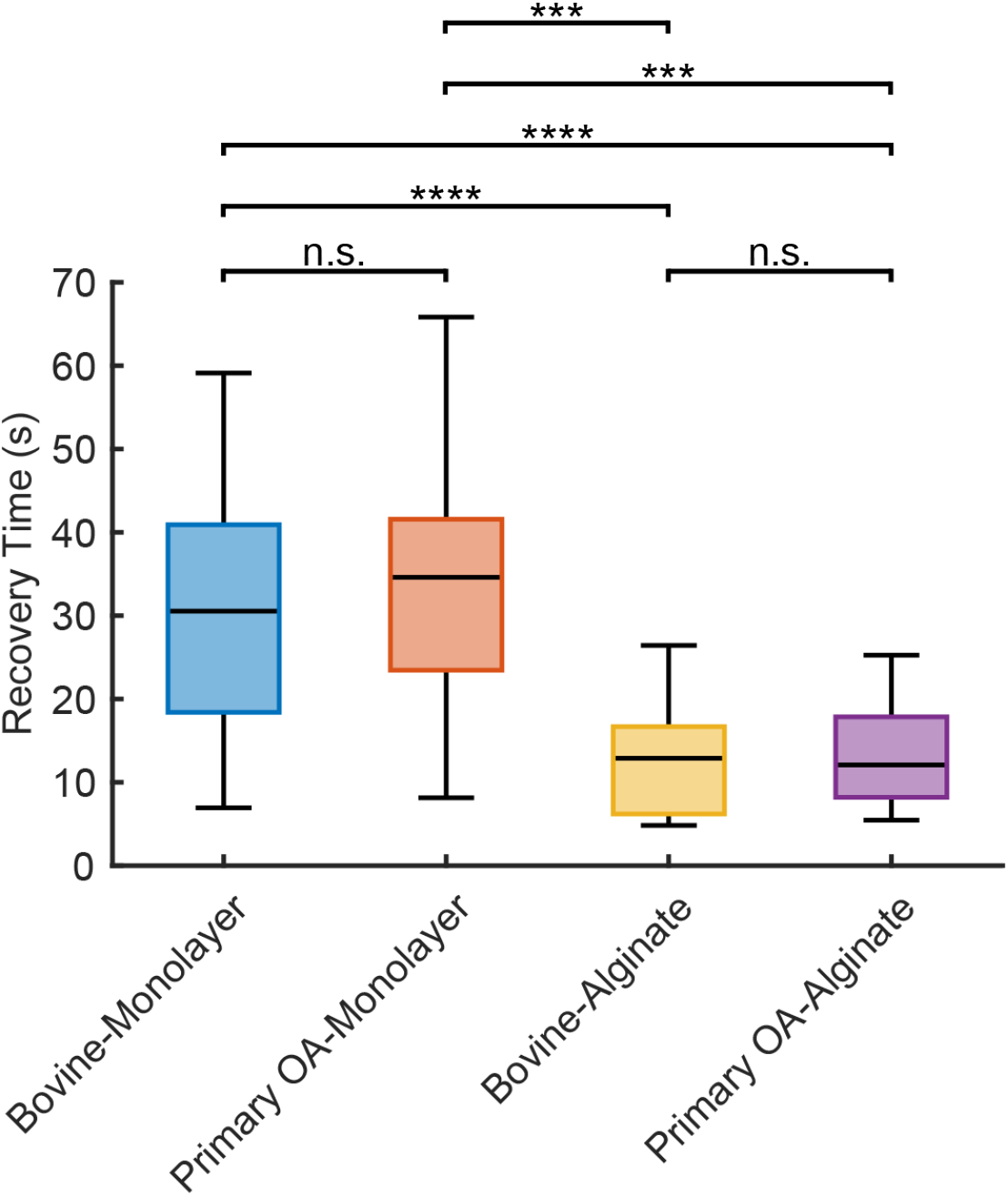
A boxplot of the viscoelastic recovery time for bovine and primary OA chondrocytes from monolayer and alginate cultures. The p-values from t-tests between groups are displayed. No statistically significant difference was found within a monolayer or alginate culture method with bovine and human cells. However, all groupings of different culture methods had a statistically significant difference of p < 0.0001 or p < 0.00001, indicated by three or four stars, respectively.

While the sample size was sufficient to distinguish between cells from the two cultures, it was limited by the short recovery time of cells from alginate cultures. Each Z-stack in the CLSM time series spans approximately 2.7 s, making it difficult to fit a recovery curve when the recovery time is short. The bovine and primary OA data sets for the alginate cultures could be doubled if the measurement frequency could be increased.

## Discussion

There was no difference in viscoelastic recovery time between bovine and primary OA chondrons which were cultured in the same manner. It has been reported that the elastic modulus of the PCM of primary chondrocytes changes with the onset of OA [26-28, 35, 41]. In this study, bovine chondrocytes were used in place of healthy primary human chondrocytes due to limited access to healthy human tissue. While the recovery time was measured in this work, it is theoretically a function of both spring constants in the Burgers model, which both contribute to the bulk elastic modulus [50]. Given that no difference in the recovery time was observed, it suggests the spring constants and, therefore, the bulk elastic modulus, were similar between monolayer bovine and primary OA chondrons.

In contrast, recovery times differed among cells from different culture methods. Monolayer and alginate bovine chondrons had different recovery times, and the same was true for monolayer and alginate primary OA chondrons. This finding is reasonable given the previously cited literature demonstrating that culture methods produce chondrocytes with different PCMs.

This study’s measurement throughput was lower than the previously reported use with chondrocytes [44], as seen by the generally low sample size. One source of this is the imaging of the PCM, and using it to define the cell border excluded cells that did not possess a nearly spherical shape due to variation in the PCM. The increased PCM on chondrocytes in this work compared to [44] added an additional challenge as the cells were more likely to stick together when in close proximity to another cell. All cells in contact with other cells were excluded from the analysis.

The overall throughput of this device and method is limited, first by the FOV of the microscopy system (0.455 mm x 0.455 mm) and second by the number of cells within that FOV. Optimizing the cell concentration to maximize the number of cells within the FOV would increase the sample size. There are several other limitations and possible improvements to this work. All experiments were conducted at room temperature (approximately 20°C), but keeping cells at 37°C or another fixed temperature would be ideal to eliminate any sources of error or variation this may introduce. Only living cells were measured; nonfluorescent and dim cells were excluded, and all cells were imaged within 90 minutes of removal from a temperature-controlled environment. Increasing the image stack height would enable visualization of the entire cell, thereby accounting for cell orientation during experiments. The stack height was limited to be less than the cell diameter to increase imaging frequency, because the current analysis assumes that chondrons deform as rigid spheroids. This simplifying assumption underestimates the linear strain of a cell compressed between glass plates, but greatly reduces measurement complexity.

## Conclusion

This work reports on the use of a previously developed 3D-printed variable-height flow cell. CLSM imaging and data fitting to a Burgers mechanical model are performed similarly; however, in this case, the cell membrane of the chondrocytes was fluorescently stained and used instead of the cytoplasm viability stain. The primary objective of this work was to determine whether monolayer and alginate cultures produced mechanically distinct chondrons. It has been previously demonstrated that 3D culture methods using alginate or hydrogels produce chondrocytes with increased PCM production compared to 2D monolayer cultures. 3D cultures allow chondrocytes to grow more naturally, as in the *in vivo* environment, where they adopt a spherical shape with a surrounding PCM. Given the prevalence of osteoarthritis and the ongoing efforts to understand its causes and develop better treatments and prevention strategies, producing chondrocytes with the same phenotype as those *in vivo* is paramount. This work provides a method for evaluating the viscoelastic properties of chondrocytes from different cultures.

The viscoelastic recovery time between bovine and primary OA chondrocytes from monolayer cultures was indistinguishable. However, the recovery time of chondrons from alginate cultures was shorter than that of chondrons from monolayer cultures, regardless of whether the chondrons were bovine or human. This suggests that the PCM, which transfers forces to the chondrocyte, is mechanically distinct from different cultures.

## CRediT authorship contribution statement

**Michael Neubauer:** Writing – Original Draft, Writing – Review & Editing, Visualization, Investigation, Methodology, Formal Analysis, Data Curation, Conceptualization.

**Priyanka Brahmachary:** Writing – Original Draft, Writing – Review & Editing, Methodology. **Ronald June:** Writing – Review & Editing, Conceptualization, Funding Acquisition. **Stephan Warnat:** Writing – Review & Editing, Conceptualization, Supervision.

## Acknowledgments

This work was supported in part by the National Institutes of Health (NIH) Grant Number R01 AR073964 and the National Science Foundation (NSF) Grant Number CMMI-2140127. Imaging was performed at the Center for Biofilm Engineering Imaging Facility at Montana State University, which is supported by funding from the National Science Foundation MRI Program (2018562), the M. J. Murdock Charitable Trust (202016116), and the US Department of Defense (77369LSRIP).

